# Bile acid supplementation improves murine pancreatitis in association with the gut microbiota

**DOI:** 10.1101/2020.02.13.948547

**Authors:** You-Dong Wan, Rui-Xue Zhu, Xin-Ting Pan

**Affiliations:** Department of Emergency Intensive Care Unit, The Affiliated Hospital of Qingdao University, Qingdao, PR China; Health Management Centre, The Affiliated Hospital of Qingdao University, Qingdao, PR China

## Abstract

Disorders of bile acids (BAs) are closely related to the development of liver and intestinal diseases, including acute pancreatitis (AP). However, the mechanism underlying the involvement of BAs in AP development remains unclear. We used intraperitoneal injection of cerulein to construct AP mouse models. These mice had significantly reduced tauroursodeoxycholic acid (TUDCA) and an imbalance of intestinal microbiota, based on 16S rDNA gene sequencing. To explore the role of AP-induced intestinal microbiota changes in the development of AP, we transplanted stool obtained from AP mice to antibiotic-treated, microbiota-depleted healthy mice. Microbiota-depleted mice presented injury to the intestinal barrier function and pancreas. Additionally, microbiota depletion reduced AP-associated pancreatic injury. This indicated that the gut microbiota may worsen AP. As TUDCA was deficient in AP mice, we gavaged AP mice with it, and evaluated subsequent expression changes in the bile acid signaling receptors farnesoid-x-receptor (FXR) and its target gene fibroblast growth factor (FGF) 15. These were downregulated, and pancreatic and intestinal barrier function injury were mitigated. Similar results were found in microbiota-depleted AP without BA treatment. However, we did not observe further downregulation of the FXR signaling pathway in microbiota-depleted AP mice given TUDCA, indicating that improvement of pancreatitis by TUDCA may be associated with gut microbiota. Our analysis of changes to the gut microbiota in AP indicated that *Lactobacilli* may be the key contributors. Taken together, our study shows that supplementation with BAs could improve bile acid-FXR-FGF15 signaling, and reduce pancreatic and intestinal injury, and that this effect may be associated with the gut microbiota.

## 1. Introduction

An important component of bile participating in fat metabolism, BAs can also act as signaling molecules by interacting with cell membranes and nuclear receptors, and play important roles in glucose and lipid metabolism and energy homeostasis [1]. Recent studies have found that dysregulated BAs are closely associated with hepatopathy, for example, steatohepatitis, hepatocellular carcinoma, and intestinal diseases such as colorectal cancer [2,3]. BAs circulating in the gastrointestinal tract are important messengers linking intestinal microecology and intestinal diseases. Specifically, intestinal microbes activate bile acid receptors, resulting in proportional and structural changes to bile acid composition, and thereby producing biological effects [4]. Two key receptors of BAs are the Takeda G-protein coupled receptor clone 5 (TGR5) and the farnesoid-X-Receptor (FXR) [1]. FXR is a ligand-dependent transcription factor, belonging to the nuclear receptor superfamily, and is mainly expressed in the liver, intestine, kidney, and adrenals. FXR activates fibroblast growth factor (FGF)19 in humans and FGF15 in mice [5]. Activation of bile acid receptors can further activate several specific signaling pathways including lipid metabolism, the immune system, signal transduction, and others[5].

Acute pancreatitis (AP) is an acute digestive disease characterized by acute upper abdominal pain, elevated serum amylase, exudation of pancreatic edema, and inflammation, with a mortality rate as high as 10–30% [6]. Recent studies indicate that intestinal microbiota participate in the occurrence and development of pancreatitis [7,8]. Our group has previously reported dramatically changed intestinal microbiota in rats with pancreatitis, and we were able to improve the intestinal microbiota imbalance of theses rats with anti-inflammatory therapy [9]. Evidence also indicates that bile acids are involved in the development of AP, but the exact mechanisms underlying this association are unclear, especially in non-biliary pancreatitis [10].

As gut microbiota regulates BA production and signaling, we assumed that AP-associated gut microbiota changes could result in alterations of bile acid profiles, activating bile acid-FXR-FGF15 signaling, and resulting in pancreas and intestinal injury. Therefore, we hypothesized that adjusting BA levels may treat AP. In order to study these questions, we undertook three principal experiments with murine models of AP. These examined microbiota composition and BA levels in AP, the effects of microbiota deletion and transplant, and the therapeutic effects of TUDCA supplementation.

## 2. Materials and methods

### 2.1. Murine AP model

This study was carried out in strict accordance with the laboratory animal management guidelines of Qingdao university, and approved by the ethics committee of Qingdao university animal experiments. Healthy male C57BL/6 mice weighing between 25 and 30 g were supplied by the Laboratory Animal Center of Qingdao University (Qingdao, China). We constructed the AP model following the method of Ding et al. [11] Briefly, mice in the AP group were given ten hourly intraperitoneal injections of a supramaximal dose of caerulein (Sigma-Aldrich, St. Louis, Missouri, USA). Lipopolysaccharide (Sigma-Aldrich, St. Louis, Missouri, USA) was administered by intraperitoneal injection immediately after the 10^th^ injection of caerulein. Mice intraperitoneally injected with saline were used as controls.

#### Experimental 1: analysis of gut microbiota and BA levels in AP

Thirty C57BL/6 mice were randomly divided into a sham-operated group (the SO group), and an AP group, with 15 mice in each group. Mice were sacrificed at 24 h after AP induction. Pancreatic and ileal tissues were harvested individually and fixed in 40 g/L formaldehyde. Samples were then embedded in paraffin and continually sectioned. We collected fresh stool samples before the mice were sacrificed and stored them at −80 °C before subjecting them to analyses of bile acid levels and microbiota (Sections 2.2 and 2.3). Blood was collected by cardiac puncture and centrifuged at 15 000 rpm for 15 min, followed by testing for trypsin and inflammatory cytokines.

#### Experimental 2: effects of gut microbiota deletion and transplant

To further explore the role intestinal microbiota plays during AP, we treated healthy mice with broad-spectrum antibiotics to delete intestinal bacteria, thus creating germ-free (GF) mice. Stool samples were collected from AP mice, then transplanted to GF mice to create fecal microbiota-transplanted (FMT) mice. Further details of the GF and FMT model mice are described in Supplementary materials 1. In brief, sixty C57BL/6 mice were randomly divided into four groups of 15 as follows: an SO group, an AP group, an GF+AP group, and an GF+FMT group. Pancreatic tissues, blood, and stool samples were collected from all mice.

#### *Experimental 3*: effects of TUDCA supplementation

In the first experiment, TUDCA was found to be reduced in AP. We therefore fed AP mice a diet of standard laboratory chow supplemented with 0.4% TUDCA (Cayman Chemicals, Michigan, USA) to explore the effect of bile acid supplementation on AP. As bile acids and their conjugated forms were identified as FXR ligands, we analyzed the bile acid-FXR-FGF15 signaling axis using RT-PCR (Section 2.5). In this experiment, sixty C57BL/6 mice were randomly divided into five groups of 12 as follows: an SO group, an AP group, an AP + TUDCA pancreatitis group, a GF pancreatitis group, and a GF+ TUDCA pancreatitis group.

### 2.2 Analysis of intestinal microbiota

Samples were analyzed by 16S rDNA gene sequencing, the details of which we have reported previously [9]. Operational taxonomic units (OTUs) that reached 97% similarity and Shannon index were used for α-diversity estimations. Nonmetric multidimensional scaling methods were conducted to visualize differences between two groups. Linear discriminant analysis was used to explore principal differences between types of bacteria. Details of our intestinal microbiota functional annotation are given in Supplementary materials 1.

### 2.3 Bile acid analysis

BA levels in feces were quantitatively measured by ultra-performance liquid chromatography triple quadrupole mass spectrometry (UPLC-TQMS) according to the following protocol. The fecal samples were extracted with methanol and the supernatant was transferred and vacuum-dried. UPLC-MS raw data obtained with negative mode were analyzed using TargetLynx applications manager version 4.1 (Waters Corp., Milford, MA) to obtain calibration equations and the quantitative concentration of each bile acid in the samples. For details, see Supplementary materials 1.

### 2.4 Histological evaluation and measurement of amylase D-lactate inflammatory cytokines

Pancreas and distal ileum samples were stained with hematoxylin and eosin (HE), and examined and scored with a published system for grading of intestinal tissue injury [12]. Amylase activity, D-lactate level, and diamine oxidase activity in serum were measured using enzyme assay kits (Shanghai Hengfei Bioscience, China). The levels of IL-1β, TNF-α, and IL-6 were measured using an ELISA kit following the manufacturer’s instructions (LMAIBio Biotech, China).

### 2.5 Real-time PCR analysis

Pancreatic tissue and intestinal mucosa scraped from the ileum were frozen in liquid nitrogen and stored at −80 °C. A standard phenol-chloroform extraction was performed to isolate total RNA from frozen tissues with Trizol reagent. Synthesis of cDNA was performed from 2μg of total RNA with a Reverse Transcription Kit (Shanghai Hengfei Bioscience, China). The real-time PCR primer sequences are listed in Supplementary materials 1.

### 2.6 Statistical analysis

Values are expressed as the mean ± SEM. Significant differences between two groups were evaluated with a two-tailed, unpaired Student’s t-test, or Mann-Whitney U test for samples that were not normally distributed. Multiple groups were analyzed by one-way or two-way ANOVA followed by Bonferoni or Dunnett’s multiple comparison test. Correlation analyses involving the gut microbiome and bile acid metabolism were performed using the nonparametric Spearman’s test. Data were subjected to statistical analysis using SPSS 15 software (SPSS, Chicago, IL); P<0.05 was considered statistically significant.

## 3. Results

### 3.1 Changes to gut microbiota and bile acid metabolism in AP mice

Acute pancreatitis was induced by intraperitoneal injections of caerulein, and was assessed based on amylase quantification and histopathological changes in pancreatic tissue. Pancreatic inflammatory cell infiltration, hemorrhage, ileal edema, and shortened villi were observed in the AP group (Figure 1A). Compared with the SO group, the levels of amylase in serum were significantly increased in the AP group (P<0.05) (Figure 1A). The above results indicated that the AP model was established successfully. To identify AP-induced changes to the composition of the gut microbiota, we conducted 16S rDNA amplicon sequencing. The Shannon diversity index and numbers of observed OTUs (α-diversity) of gut microbiota were remarkably decreased after AP induction (Figure 1B). Our nonmetric multidimensional scaling method showed that the gut microbiota composition was substantially reshaped in the AP group (Figure 1C). Linear discrimination analysis coupled with effect size analysis revealed significant increases of *Lactobacillus* and *Escherichia-Shigella* and substantial reductions of *Roseburia, Ruminococcaceae_NK4A214_group, norank_f_Bacteroidales_ S24-7_group*, and *unclassified_f_Peptostreptococcaceae* in AP compared with SO (Figure 1D). To characterize the functional alterations of the gut microbiota in AP, the relative abundances of KEGG pathways predicted by PICRUSt were calculated based on the 16S rRNA sequencing data. Multiple KEGG categories were disturbed in AP compared to SO. There was significant enhancement of the pathways for infectious diseases, metabolism of terpenoids and polyketides, immune system diseases, and signal transduction, and significant weakening of genetic information processing, the circulatory system, transcription, and digestive system pathways (Supplementary material 2A).

**Figure 1.**
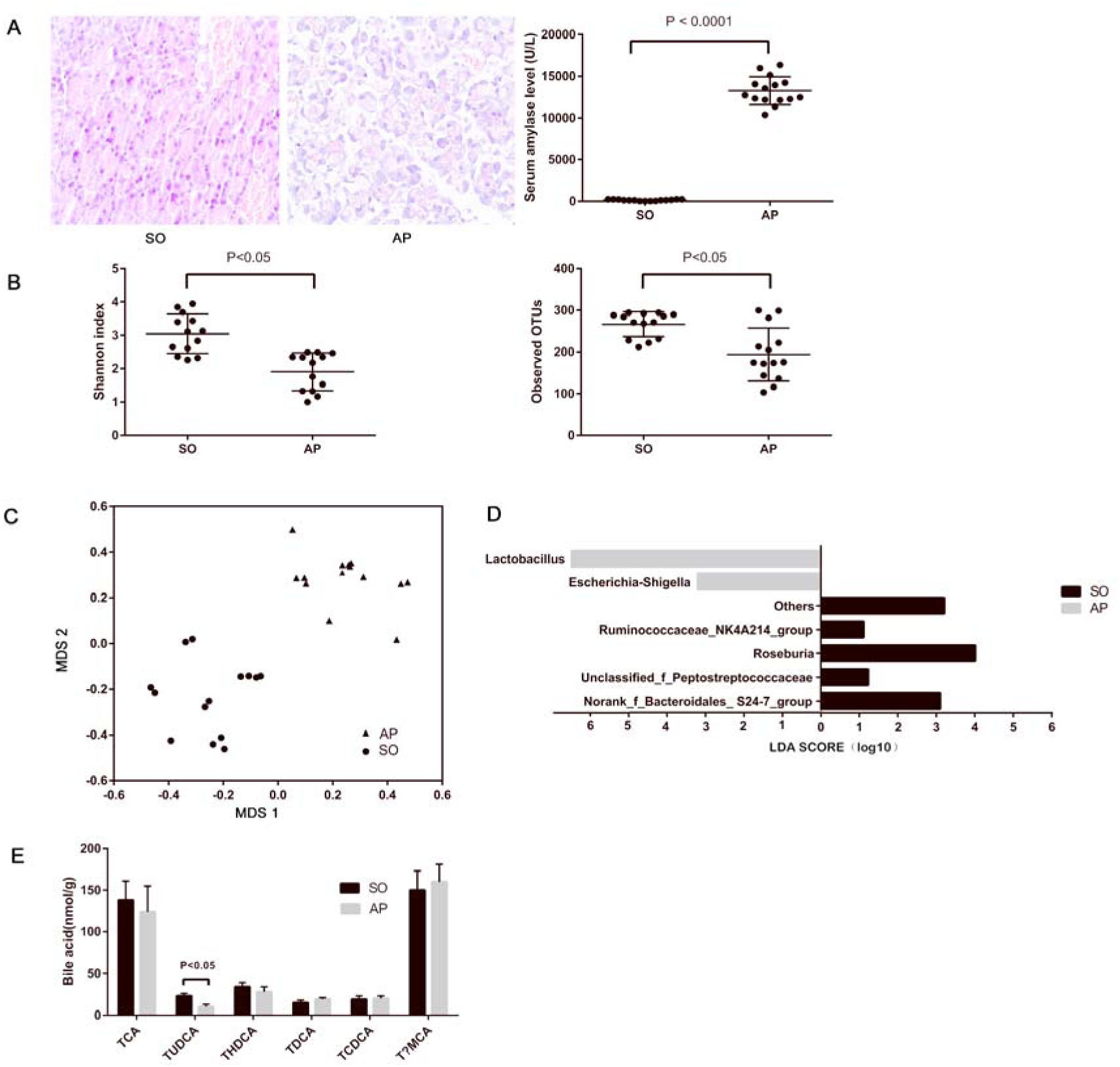
Gut microbiota and bile acid metabolism changes in AP mice. (A) Histological analysis of the pancreas and levels of serum amylase in sham-operated (SO) and acute pancreatitis (AP). (B) The Shannon diversity index and numbers of observed OTUs (α-diversity) are different between AP and SO. (C) Nonmetric multidimensional scaling method showing a definite shift in gut microbiota composition between AP and SO. The horizontal axis represents the first dimension and the vertical axis represents the second dimension. (D) Linear discriminant analysis scores for the bacterial taxa differentially abundant between acute pancreatitis and the sham-operated group. Only the taxa having a p value < 0.01 and LDA > 2.0 were shown. (E) Stool bile acid levels in the SO and AP groups.

We used UPLC-TQMS metabolite profiling to quantitate bile acid levels in the stool. The level of TUDCA was significantly decreased in the AP group (Figure 1E). Total bile acid levels remained unchanged, but noticeable decreases in ratios of conjugated to unconjugated BAs were observed. There was no difference in the ratio of 12 α-OH to non-12 α-OH bile acids (Supplementary materials 2B). Altogether, these data suggested that AP changed the gut microbiota and bile acid metabolism in mice.

### 3.2 AP-associated changes to the gut microbiota aggravate AP

To explore the role of AP-induced intestinal microbiota changes in the development of AP, we used four groups of mice: SO, AP, GF+AP, and GF mice given fecal transplant from AP mice (GF+FMT). Pancreatic histopathological changes were mitigated in the GF+AP group compared to the AP group (Figure 2A). Based on plasma D-lactate and diamine oxidase levels, intestinal barrier function injury was mitigated in GF+AP (Figure 2A). Plasma levels of TNF-α, IL-1β, and IL-6 were significantly decreased in the GF+AP group (Figure 2B). Interestingly, in the GF+FMT group, mildly aggravated histology, intestinal barrier function, and plasma inflammation were found. Altogether, these data suggested that AP gut microbiota aggravate AP.

**Figure 2.**
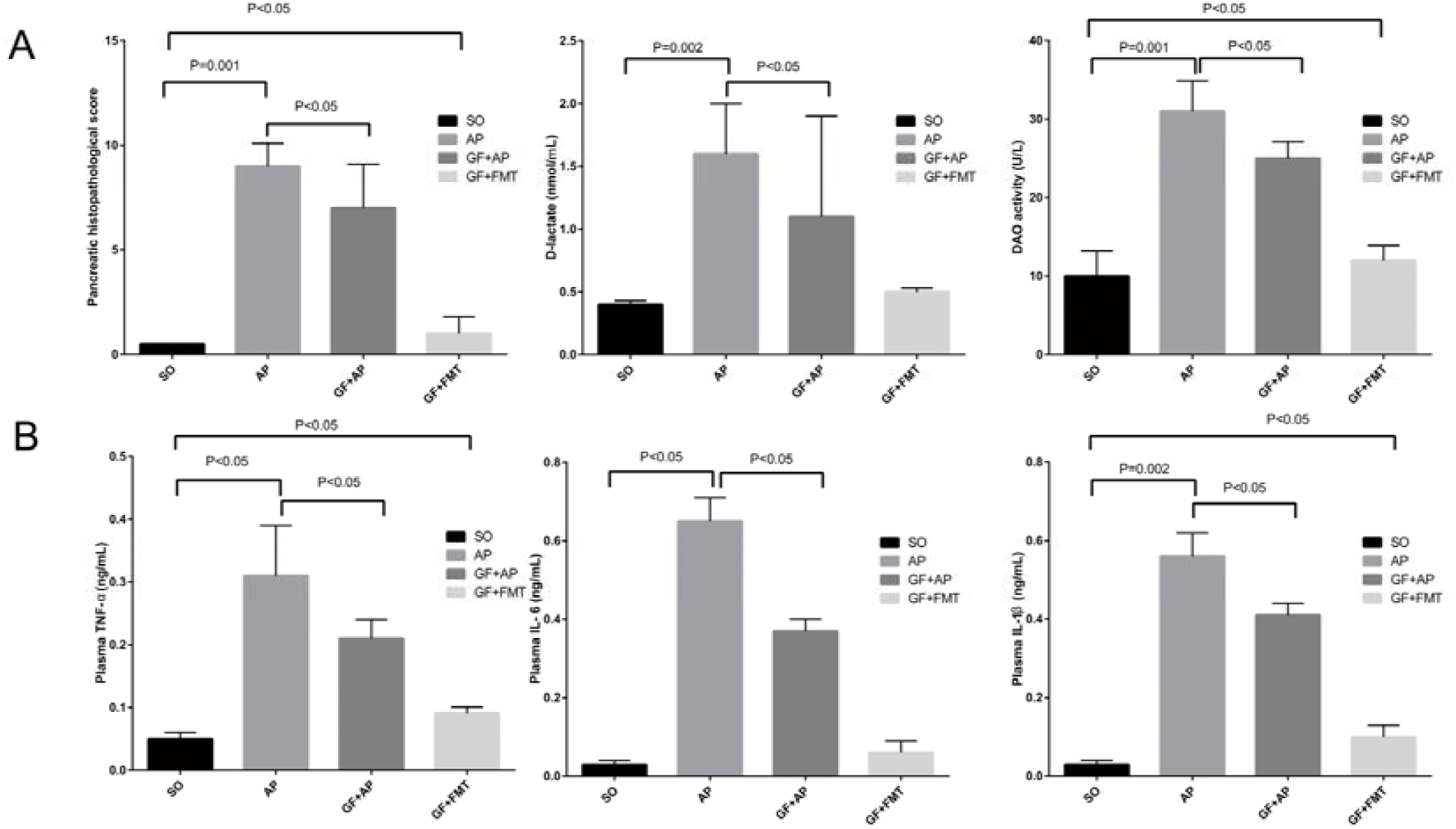
Acute pancreatitis gut microbiota worsen pancreatitis. (A) Changes to pancreatic histopathology, plasma D-lactate, and diamine oxidase between sham-operated (SO), acute pancreatitis (AP), germ-free (GF), and fecal microbiota transplantation (FMT) mice. (B) Changes to levels of intestinal inflammatory cytokines (tumor necrosis factor α, IL-1β, and IL-6).

### 3.3 TUDCA has gut microbiome-associated therapeutic effects on AP

To explore the effect of bile acid supplementation, we gave AP mice TUDCA, in view of its observed reduction in AP. This resulted in significantly decreased plasma D-lactate, diamine oxidase, TNF-α, IL-1β, and IL-6 levels, indicating improved intestinal barrier function and ameliorated systemic inflammatory response during AP. (Figure 3A, and Supplementary material 3). Although bile acids and their conjugated forms have been identified as FXR or TGR5 ligands, a previous study demonstrated that TUDCA has no effects on TGR5 activity [13]. However, activation of the FXR-small heterodimer partner (SHP) inhibits bile acid synthesis. Therefore, we evaluated expression of FXR target gene mRNAs including FXR, FGF15, and SHP. We found that the intestinal FXR-FGF15 axis was downregulated with TUDCA supplementation in AP mice (Figure 3B). To investigate whether the gut microbiota were involved in inhibition of intestinal FXR signaling by bile acid, GF mice were used. These mice showed downregulation of the FXR-FGF15 axis during AP, but further inhibition of FXR signaling was not observed with the addition of TUDCA (Figure 3C). These results revealed the involvement of the gut microbiota in inhibiting intestinal FXR signaling by TUDCA. Correlating bacterial general with TUDCA, we found positive associations for *Lactobacillus* (P<0.05), and some other AP-enriched bacteria including *Ruminococcus 2, Prevotellaceae UCG-003*, and *Ruminococcus 1* (Figure 3D). Altogether, these data suggested that TUDCA has therapeutic effects on AP associated with the gut microbiota.

**Figure 3.**
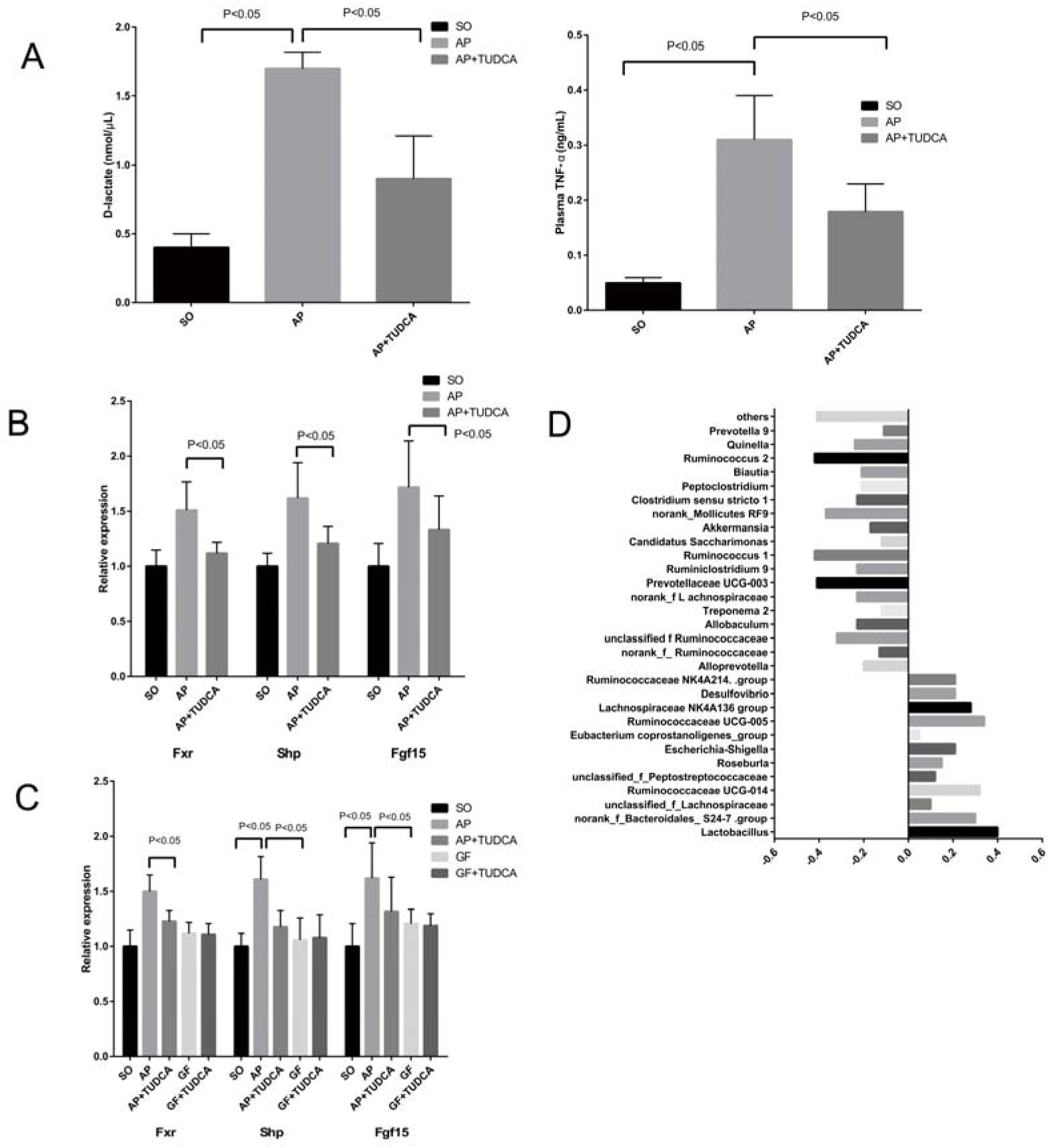
Tauroursodeoxycholic acid (TUDCA) has therapeutic effects on acute pancreatitis associated with gut microbiota. (A) Changes of D-lactate and tumor necrosis factor α levels between the three groups. (B) Relative expression of intestinal FXR mRNA and its target gene mRNAs in mice. (C) Relative expression of intestinal Fxr, Shp, and Fgf15 mRNAs in mice. (D) Spearman correlations between the most abundant genera and TUDCA.

## 4. Discussion

The invasion of bile acid into the pancreatic duct, caused by bile reflux, has traditionally been considered part of the pathogenesis of acute biliary pancreatitis. However, the role and mechanism of bile acid in non-biliary pancreatitis is still unclear. In this study, the level of TUDCA was reduced in mice with non-biliary pancreatitis. With TUDCA supplementation, the AP mice showed improved pancreatic and intestinal injury and decreased systemic inflammatory response. Further study indicated that a possible underlying mechanism was downregulation of bile acid-FXR-FGF15 signaling, the upregulation of which is associated with pancreatitis.

TUDCA is a nontoxic taurine conjugate form of ursodeoxycholic acid, which is an endogenously produced hydrophilic bile acid. Several studies have shown its potential for treating liver diseases [16], and possible mechanisms underlying this include prevention of cell death by stabilization of the cell membranes, inhibition of apoptosis, and upregulation of survival pathways[17]. However, to our knowledge, the effects of TUDCA on AP have not been investigated previously. At present, the main AP treatments include fluid resuscitation, nutritional support, and infection prevention [18]. The present study indicates the potential of TUDCA as a new horizon for the treatment of AP.

In view of evidence that the gut microbiota regulate bile acid production and signaling [4], we also investigated their relationship with AP and BAs. Our fecal transplantation experiment showed that the gut microbiota from AP mice were harmful to the intestinal function of healthy mice. Moreover, although deletion of gut microbiota and supplementation with TUDCA were both useful for the treatment of AP, our third experiment showed that deletion of gut microbiota plus TUDCA supplementation resulted in no additional therapeutic effect, with gene expression analysis revealing that gut microbiota may participate in inhibition of intestinal FXR signaling via TUDCA.

Several studies have explored the association between gut microbiota and AP by intestinal gene sequencing. Zhu et al. [14] found that increased capacity for bacterial invasion of epithelial cells in AP correlated closely with the abundance of *Escherichia-Shigella* in fecal samples from 165 adults. Zheng et al.[15] showed that commensal *E. coli* MG1655 increases TLR4/MyD88/p38 MAPK and ERS signaling-induced intestinal epithelial injury and aggravates AP in rats. These authors inferred gut microbiota dysbiosis in AP, and tried to explain it from different perspectives. Similar to their results, our study also found dysbiosis of the gut microbiota in AP, and the altered BA metabolism we observed is a novel potential mechanism.

The initial and major sites of injury in acute pancreatitis are the pancreas and intestines. Bile acids are critical components of the gastrointestinal tract that link the gut microbiota to hepatic and intestinal metabolism. The ability of the gut microbiota to biotransform intestinal bile acids into their unconjugated forms is central to the metabolic homeostasis of the gastrointestinal tract [4].The main bacterial genera of gut microbiota involved in bile acid metabolism include *Bacteroides, Clostridium, Lactobacillus, Bifidobacterium*, and *Listeria* [19]. Pancreatitis can damage the intestinal micro-environment, and thus change BA metabolism. Disruption of bile acid-microbiota crosstalk can promote inflammation, organ injury, and gastrointestinal disease phenotype, which can contribute to the development of gastrointestinal cancers, including colorectal cancer and hepatocellular carcinoma [20]. In our study, disruption of bile acid-microbiota crosstalk manifested as decreases in the variety of bacterial species and their proportions, and functional alterations of the gut microbiota were reflected by decreased ratios of conjugated to unconjugated bile acids, and TUDCA deficiency. These changes were associated with further injury to the pancreas and intestines. In a study by Sun et al. [13], TUDCA was confirmed as an FXR antagonist in vitro and in vivo, and our study yielded similar results.

Several limitations of this study should be addressed. Firstly, the effect of BAs on changes to the FXR-FGF 15 axis we observed need further verification with intestinal-specific Fxr knockout mice and floxed control mice, which is an aim of our future work. Secondly, we showed that *Lactobacillus* was positively correlated with TUDCA, but the relationships among *Lactobacillus*, BAs, and the FXR-FGF 15 axis during AP are unclear and deserve further study. Thirdly, we found that a significant number of bacterial species were unidentifiable because of technological limitations, and with advances in sequencing technology more gut bacteria relevant to pancreatitis may be detected.

In conclusion, BA supplementation could improve bile acid-FXR-FGF15 signaling, and reduce pancreatic and intestinal injury in AP, and this effect may be associated with the gut microbiota. It is possible that TUDCA may be a promising therapeutic treatment for AP in clinical practice.

## Acknowledgments

This study was supported by the National Natural Science Foundation of China (Grant No. 81870440)

## References

[1] Martinot E, Sedes L, Baptissart M, Lobaccaro JM, Caira F, et al. (2017) Bile acids and their receptors. Mol Aspects Med 56: 2–9.

[2] Arab JP, Karpen SJ, Dawson PA, Arrese M, Trauner M (2017) Bile acids and nonalcoholic fatty liver disease: Molecular insights and therapeutic perspectives. Hepatology 65: 350–362.

[3] Joyce SA, Gahan CG (2017) Disease-Associated Changes in Bile Acid Profiles and Links to Altered Gut Microbiota. Dig Dis 35: 169–177.

[4] Ramirez-Perez O, Cruz-Ramon V, Chinchilla-Lopez P, Mendez-Sanchez N (2017) The Role of the Gut Microbiota in Bile Acid Metabolism. Ann Hepatol 16: s15–s20.

[5] Kliewer SA, Mangelsdorf DJ (2015) Bile Acids as Hormones: The FXR-FGF15/19 Pathway. Dig Dis 33: 327–331.

[6] Tenner S, Baillie J, DeWitt J, Vege SS (2013) American College of Gastroenterology guideline: management of acute pancreatitis. Am J Gastroenterol 108: 1400–1415, 1416.

[7] Tan C, Ling Z, Huang Y, Cao Y, Liu Q, et al. (2015) Dysbiosis of Intestinal Microbiota Associated With Inflammation Involved in the Progression of Acute Pancreatitis. Pancreas 44: 868–875.

[8] Signoretti M, Roggiolani R, Stornello C, Delle FG, Capurso G (2017) Gut microbiota and pancreatic diseases. Minerva Gastroenterol Dietol 63: 399–410.

[9] Wan YD, Zhu RX, Bian ZZ, Pan XT (2019) Improvement of Gut Microbiota by Inhibition of P38 Mitogen-Activated Protein Kinase (MAPK) Signaling Pathway in Rats with Severe Acute Pancreatitis. Med Sci Monit 25: 4609–4616.

[10] Hegyi P, Maleth J, Walters JR, Hofmann AF, Keely SJ (2018) Guts and Gall: Bile Acids in Regulation of Intestinal Epithelial Function in Health and Disease. Physiol Rev 98: 1983–2023.

[11] Ding SP, Li JC, Jin C (2003) A mouse model of severe acute pancreatitis induced with caerulein and lipopolysaccharide. World J Gastroenterol 9: 584–589.

[12] Chiu CJ, McArdle AH, Brown R, Scott HJ, Gurd FN (1970) Intestinal mucosal lesion in low-flow states. I. A morphological, hemodynamic, and metabolic reappraisal. Arch Surg 101: 478–483.

[13] Sun L, Xie C, Wang G, Wu Y, Wu Q, et al. (2018) Gut microbiota and intestinal FXR mediate the clinical benefits of metformin. Nat Med 24: 1919–1929.

[14] Zhu Y, He C, Li X, Cai Y, Hu J, et al. (2019) Gut microbiota dysbiosis worsens the severity of acute pancreatitis in patients and mice. J Gastroenterol 54: 347–358.

[15] Zheng J, Lou L, Fan J, Huang C, Mei Q, et al. (2019) Commensal Escherichia coli Aggravates Acute Necrotizing Pancreatitis through Targeting of Intestinal Epithelial Cells. Appl Environ Microbiol 85.

[16] Carey EJ, Ali AH, Lindor KD (2015) Primary biliary cirrhosis. Lancet 386: 1565–1575.

[17] Schoemaker MH, Conde DLRL, Buist-Homan M, Vrenken TE, Havinga R, et al. (2004) Tauroursodeoxycholic acid protects rat hepatocytes from bile acid-induced apoptosis via activation of survival pathways. Hepatology 39: 1563–1573.

[18] Vege SS, DiMagno MJ, Forsmark CE, Martel M, Barkun AN (2018) Initial Medical Treatment of Acute Pancreatitis: American Gastroenterological Association Institute Technical Review. Gastroenterology 154: 1103–1139.

[19] Gerard P (2013) Metabolism of cholesterol and bile acids by the gut microbiota. Pathogens 3: 14–24.

[20] Yoshimoto S, Loo TM, Atarashi K, Kanda H, Sato S, et al. (2013) Obesity-induced gut microbial metabolite promotes liver cancer through senescence secretome. Nature 499: 97–101.

